# AxonFinder: Automated segmentation of tumor innervating neuronal fibers

**DOI:** 10.1101/2024.09.03.611089

**Authors:** Kaoutar Ait-Ahmad, Cigdem Ak, Guillaume Thibault, Young Hwan Chang, Sebnem Ece Eksi

## Abstract

Neurosignaling is increasingly recognized as a critical factor in cancer progression, where neuronal innervation of primary tumors contributes to the disease’s advancement. This study focuses on segmenting individual axons within the prostate tumor microenvironment, which have been challenging to detect and analyze due to their irregular morphologies. We present a novel deep learning-based approach for the automated segmentation of axons, AxonFinder, leveraging a U-Net model with a ResNet-101 encoder, based on a multiplexed imaging approach. Utilizing a dataset of whole-slide images from low-, intermediate-, and high-risk prostate cancer patients, we manually annotated axons to train our model, achieving significant accuracy in detecting axonal structures that were previously hard to segment. Our analysis includes a comprehensive assessment of axon density and morphological features across different CAPRA-S prostate cancer risk categories, providing insights into the correlation between tumor innervation and cancer progression. Our paper suggests the potential utility of neuronal markers in the prognostic assessment of prostate cancer in aiding the pathologist’s assessment of tumor sections and advancing our understanding of neurosignaling in the tumor microenvironment.

## INTRODUCTION

Neurosignaling has emerged as a hallmark feature in cancer research ^1^, highlighting the importance of understanding the intricate relationship between neuronal innervation of primary tumors and cancer progression ^2^. The dynamic interplay between cancer cells and their microenvironment is crucial for understanding tumor progression for effective therapies. Recent research has underscored neurosignaling as a significant factor in this interplay, particularly through its role in promoting cancer cell proliferation, migration, and invasion ^3–7^. The neuro-oncological interface is complex, involving multiple signaling pathways that contribute to the cancer’s ability to grow and metastasize. ^8,9^ has illustrated how adrenergic nerves can foster a microenvironment conducive to cancer progression in models of prostate cancer, providing insights into how nervous system elements can directly influence patients’ outcomes^10,11^.

In the tumor microenvironment, two primary types of neuronal innervation are observed ^12,13^: neuronal bundles and neuronal fibers, i.e., axons. Neuronal bundles, comprised of clusters of nerve fibers within and around tumors, have historically received significant attention due to their macroscopic visibility. Perineural invasion, characterized by cancer cells infiltrating these neuronal bundles, has become a landmark phenotype for pathologists grading solid tumors in tissue sections ^14,15^. An intriguing gap in this research landscape is the absence of studies focused on segmenting and quantifying neuronal fibers or individual axons in tissue sections.

Neuronal fibers have been reported to impact cancer progression in prostate ^16^, head and neck ^17^, and pancreatic ^18^ cancers, stimulating the proliferation of cancer cells as a result of signaling from axon terminals. However, the staining and the unique unconventional shapes ^19^ of these individual axons have posed challenges that hindered their comprehensive investigation.

In response to these challenges, our research leverages imaging technologies and machine learning techniques to better understand and quantify the role of neuronal fibers in tumors. By employing cycIF imaging, we are able to simultaneously visualize multiple biomarkers, enhancing our understanding of the spatial and molecular context of axonal interactions within tumors. Here, we develop a deep learning model to do semantic segmentation for the axons in the prostate tumor microenvironment of patient samples ^20 21^. Contrary to studies that concentrate on neuronal bundles ^22 23^, our model provides an automated process for segmenting neuronal fibers in tissue sections based on the expression of two markers that are commonly expressed in mature axons: TUBB3 and NCAM1. To generate an efficient deep-learning model, we used an immunofluorescent dataset comprising of whole-slide images (WSI) from 17 patients obtained from OHSU’s Biolibrary. 997 axons in five tissue sections were manually annotated using the Napari software ^24^. Our results show that axonal features can provide information on prostate cancer staging. Automated segmentation of tumor-innervating axons can be used to investigate the specific impact of neuronal innervation on cancer progression ^25^ paving the way for novel insights into cancer biology.

The potential of these technological advancements extends well into clinical applications. By providing a more nuanced understanding of tumor innervation, our study may lead to novel diagnostic and prognostic tools. For instance, the density and morphology of axonal networks within tumor tissues could serve as biomarkers for aggressive cancer phenotypes, guiding treatment decisions and potentially indicating targets for therapies aimed at disrupting neuro-cancer interactions ^25^. Moreover, our model provides a computational tool to study other diseases where neurosignaling plays a critical role, such as neurodegenerative disorders ^26^. The methodology developed through this research might also be applicable in other contexts where complex tissue environments need to be analyzed, paving the way for cross-disciplinary applications in biomedical research.

## RESULTS

### The development of a U-Net axon segmentation model AxonFinder

Our U-Net model performance leveraging TensorFlow 2 and Keras 2 ^27,28^ to perform neuron segmentation on cycIF images was assessed using IoU and F1-Score as in the previous sections. Throughout the training, the models exhibited a rapid decrease in loss during training, suggesting that our model generalizes well and is not prone to overfitting. The IoU and F1-Score metrics for both training and validation datasets showed substantial improvement, with the metrics plateauing at high levels, reflecting robust segmentation capability. Figure 1 summarizes the workflow and the performance metrics for each scenario, the validation F1-Score reaches 94.81% in the case of the combined dataset (TUBB3+NCAM1), and the IoU achieves a similar level of performance of 90.78 %. On the other hand, the TUBB3 dataset shows a slightly lower IoU of 89.79% and an F1-Score of 94.18%. This convergence is a clear indication that the model has learned the intricate patterns within the cycIF images. With the improved performance observed, the merged dataset of TUBB3 and NCAM1 was utilized for the remainder of the analysis.

The results in Figure 2 underscore the efficacy of the model in the automated segmentation of axons. The first row of images are three samples from the TUBB3+NCAM1 dataset. Thus, in the second row, the green, red, and yellow colors represent the true positive, the true negative, and the overlap between these two, respectively. A strong concordance was observed between the ground truth and the predicted segmentation masks produced. Given the challenges associated with the manual segmentation of axons, due to their inconsistent features (e.g. shape and signal intensity), our model has been instrumental in identifying new axons that may have been previously overlooked. Finally, we employed the aforementioned tiling strategy, whereby each tile underwent a separate segmentation procedure using our U-Net model to delineate the neuronal structures. Subsequently, we stitched all the tiles together to reconstruct the original tissue.

**Figure 1.** Schematic representation showing the training of AxonFinder. The input image consists of a combination of TUBB3 (red) and NCAM1 (green), which are tiled across the entire tissue section (left). The images were manually annotated, and binary masks were generated. The model uses a ResNet101 encoder network, where each layer is structured with 3×3 convolutional operations. The model performance was evaluated using IoU and F1-Scores for both TUBB3 (single channel) and TUBB3 and NCAM1 (combined) segmentation results.

**Figure 2.** AxonFinder achieves segmentation in whole-section patient samples. (A) Examples of NCAM1+TUBB3 images that correspond to different patients in the test dataset. (B) The green, red and yellow colors illustrate the true positive, the true negative and the overlap between the ground truth and the prediction, respectively.

### The density of axonal fibers across Gleason grades

Upon completion of the axon segmentation and the extraction of axon features of all samples, we proceeded to conduct a thorough statistical evaluation of the data. Figure 3 presents a comprehensive view of axon density across different prostate cancer risk categories delineated by CAPRA-S and across different Gleason scores. Figure 3A shows two boxplots representing axon density within the whole-tissue area. On average, low-risk patients had 58 *axons*/*mm*^*2*^, while intermediate-risk patients had 37*axons*/*mm*^*2*^ and high-risk patients had 5 *axons*/*mm*^*2*^. The decrease in axon density from low-risk to high-risk could be attributed to the typically reduced stromal region in high-risk patients. To address this possibility, we calculated axon density using the area marked by *α*SMA marker, which marks the stromal regions ^29^, as shown in Figure 3B. Our results show a similar progressive decline in median axon density with increasing risk, suggesting that the decrease in axonal density is independent of the reduction in stromal area in high-risk patients. ANOVA analysis ^30^ showed that the density of axons per tissue and CARPA-S risk ^31^ or Gleason pattern group are significantly associated with high-risk patients as compared to intermediate- and low-risk (p < 0.05).

**Figure 3.** High-risk prostate cancer patients show a decrease in axonal density. (A) Illustration of axon density with respect to CARPA-S risk (top) and primary Gleason pattern (bottom). (B) Illustration of axon density using only the stromal areas (αSMA positive regions) with respect to CARPA-S risk (top) and primary Gleason pattern (bottom).

### Moran’s I and Ripley’s L for NCAM1 and TUBB3 marker expression

To gain a deeper understanding of the spatial distribution of segmented axons in our data set, we calculated Moran’s I ^32 33^ values with respect to CARPA-S risk scores (Supplementary Figure 1A and Supplementary Figure 1D). We used the mean intensity of TUBB3 and NCAM1 markers within the axon segmentation masks to assess spatial autocorrelation. We labeled axons ‘high’ and ‘low’ based on the expression levels of TUBB3 or NCAM1 (Supplementary Figure 1B and Supplementary Figure 1E). We then performed Ripley’s L ^34^ to calculate the spatial clustering of ‘high’ or ‘low’ axons (Supplementary Figure 1C and Supplementary Figure 1F). We found that 13 out of 17 samples showed significant results for TUBB3, revealing spatial clustering of axons expressing high levels of TUBB3 protein. Similarly, axons expressing high NCAM1 marker expression showed significant spatial autocorrelation in the same samples (Supplementary Figure 1C). These clusters imply a correlation where high TUBB3 expression is indicative of low NCAM1 expression. The trend was particularly observed in a subset of patients, with 4 out of 17 exhibiting the highest axon densities. In such dense axonal tissues, Ripley’s L analysis further corroborated the presence of more clustered axons based on the level of marker expression.

### The morphological features of axons based on patient-risk

To further investigate tumor innervation using our axon segmentation mask, we developed an analytical framework designed to establish benchmarks for the morphological characteristics of axons (Supplementary Figure 2A). The axon traits evaluated by this model are their size, shape, and branching complexity. Using a Gaussian Mixture Model (GMM), we stratified these features into specific morphological categories: Large vs. Small, Elongated vs. Circular, and Branched vs. Unbranched (Supplementary Figure 3).

The distribution graphs in Supplementary Figure 2B display a comparison of morphological features across the three risk categories. Our results show a consistency of median values for each morphological feature for low-, intermediate-, and high-risk patients. This lack of significant deviation in the medians suggests that the morphological characteristics of axons, as identified by our model, are not influenced by the varied risk categories (Supplementary Figure 2C). Such an observation is crucial, as it emphasizes the DL model’s precision in the extraction and analysis of distinct axonal features, bolstering confidence in the model’s application for evaluating neuronal innervation.

The CAPRA-S-based clusters identified in our analysis do not correlate with the axons’ features. This finding prompted us to pursue additional experimental inquiry, as illustrated in Supplementary Figure 4. We used K-means clustering to categorize the same axonal features into distinct groups. We used the elbow method to determine the optimal number of clusters, resulting in three distinct clusters ^35^. The heatmap shows these clusters based on area, solidity, and eccentricity (Figure 4A). Our clustering results identified axons that are large, branched, and elongated in Cluster 1; small, unbranched, and elongated in Cluster 2; small, unbranched, and circular in Cluster 3 (Figure 4B). We did not observe a significant correlation between risk and morphological features (Figure 4A), supporting our results in Supplementary Figure 2.

**Figure 4.** Clustering of axons based on morphological features and marker expression profiles. (A) Heatmap showing normalized axon feature values for the three clusters identified in our dataset. (B) Percentage distribution of axons characterized by different features in three distinct clusters. Cluster 1 reveals large, branched, and elongated axons; Cluster 2 shows small, unbranched, and elongated axons; Cluster 3 identified small, unbranched, and circular axons in the prostate tumor microenvironment of patients.

## DISCUSSION

The identification of tumor innervation as a fundamental hallmark of cancer has highlighted the need for automated tools capable of segmenting axons in patient samples. Addressing this gap, our study introduces a novel Res-U-Net based algorithm designed to segment axons in patient samples using two key axonal markers, TUBB3 and NCAM1. AxonFinder can be utilized to capture neuronal fibers in FFPE tissue sections, providing high-resolution proteomic content for innervated organs.

Our findings reveal a significant correlation between prostate cancer stage and axonal density, emphasizing the necessity for further investigation in larger cohorts to validate this association. Notably, high-risk patients within our cohort exhibited smaller stromal regions, correlating with disease progression. Despite efforts to normalize axon density using αSMA regions, the inclusion of an expanded sample size may clarify additional significant associations between disease grade and axon density. Moreover, our study identifies a notable phenomenon wherein axons expressing high levels of TUBB3 or NCAM1 demonstrate significant spatial autocorrelation in a subset of patients, underscoring the proteomic variability inherent in tumor-innervating axons. Future research endeavors will delineate the spatial interactions between distinct axon subtypes, thereby enriching our understanding of tumor microenvironment dynamics. Furthermore, leveraging morphological features, we recognize three distinct clusters of axons within the prostate tumor microenvironment, although no visible differences across grades were observed. Nevertheless, potential correlations between protein expression and morphological features of tumor-innervating axons demand exploration in multiplex imaging datasets.

The segmentation pipeline developed in this study holds promise not only for facilitating the study of cancer-neuron interactions among cancer biologists employing spatial imaging but also for its potential impact on elucidating the underlying biology of neurodegenerative diseases such as Alzheimer’s or Parkinson’s. By enabling the extraction of proteomic data from axons through image segmentation, this pipeline has the potential to accelerate transformative advancements in disease pathology.

### Limitations of Study

The limitations of our study that focused on automated segmentation of tumor-innervating neuronal fibers using advanced imaging and deep learning techniques stem from several factors that constrain the general inferences and applications of this research. These include the sample size and marker diversity, which may affect the generalizability of the findings based on tumor stage. Many FFPE sections are available to further investigate the results from this segmentation model, however, a dual TUBB3 and NCAM1 staining is not readily available for many public data sets. The high computational demand for training on large, high-resolution datasets can restrict replication and extension by researchers with fewer resources. Biological variability, such as differences in axon morphology across various diseases and distinct tumor microenvironments, can challenge the model’s generalizability. Translating findings into clinical practice involves additional challenges like validation across larger cohorts and integration with existing diagnostic workflows.

## Acknowledgments

We are grateful to the Biolibrary and the Histopathology Shared Resource (HSR) at OHSU for providing patient samples, specifically Aletha Letsch and Cheyenne Martin. The Biolibrary and HSR were supported by NIH grants P30 CA069533 and P30 CA069533 13S5 through the Knight Cancer Institute. Our research complies with all relevant ethical regulations of the Knight Cancer Institute at OHSU. Samples from patients undergoing radical prostatectomy were collected with informed consent through Biolibrary under IRB#4918. Participants did not receive compensation. We thank Stefanie Kaech Petrie and her team at the Advanced Light Microscopy Core at OHSU for their technical help. This project was supported by funding (CEDAR3410918) from the Cancer Early Detection Advanced Research Centre at Oregon Health & Science University, Knight Cancer Institute (S.E.E.).

## Author Contributions

KAA: Data processing, Software, Visualization, Formal Analysis, Methodology, Writing-original draft, Writing-review and editing. CA: Software, Formal Analysis, Methodology, Writing-review and editing. GT: Software, Investigation, Methodology, Supervision, Writing-review and editing. YHC: Conceptualization, Formal Analysis, Investigation, Methodology, Supervision, Writing– review and editing. SEE: Data generation, Conceptualization, Formal Analysis, Investigation, Methodology, Supervision, Writing–review and editing.

## STAR*METHODS

## KEY RESOURCES TABLE

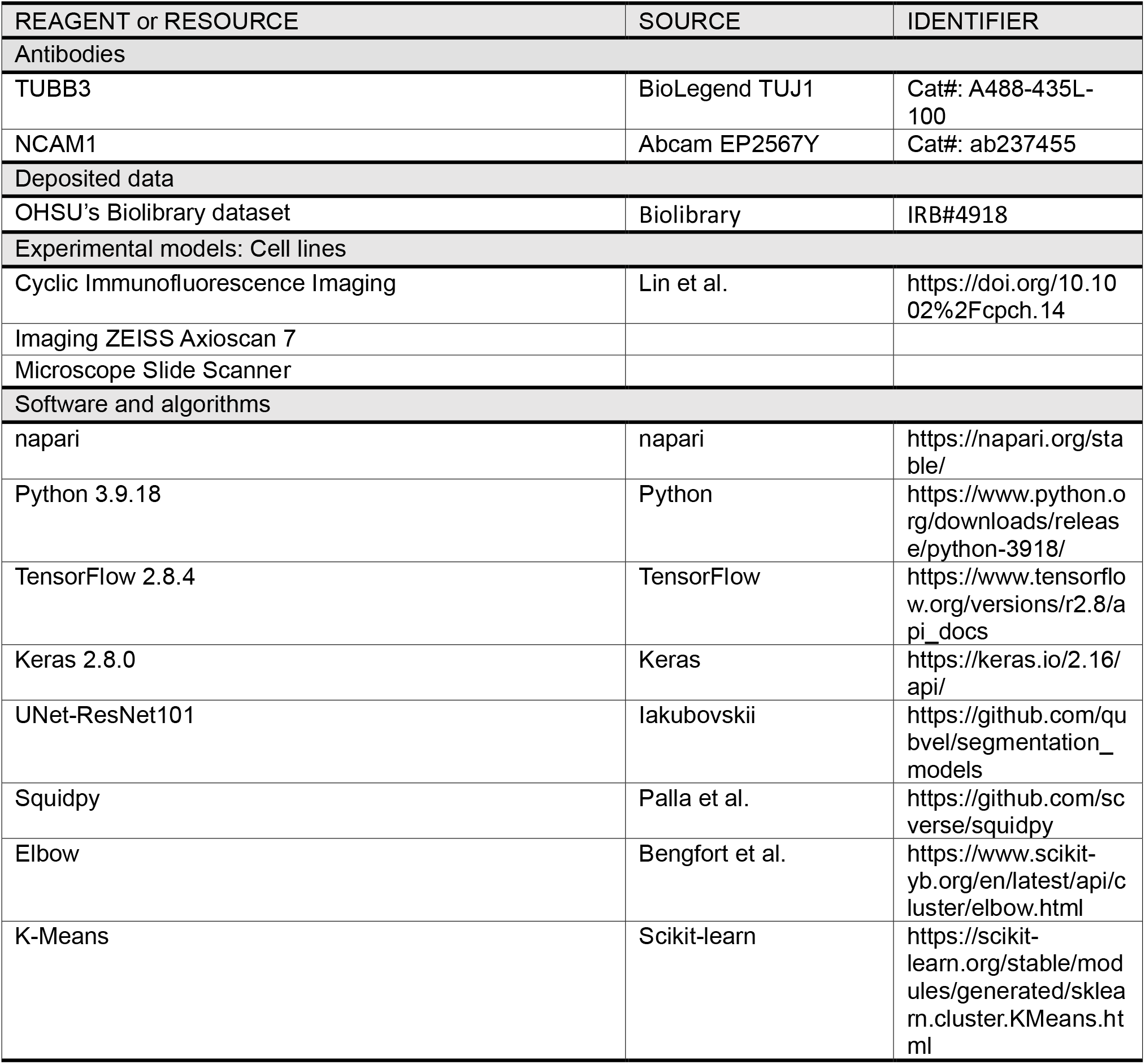

## RESOURCE AVAILABILITY

### Lead contact

Further information and requests for resources should be directed to and will be fulfilled by the lead contact, Sebnem Ece Eksi.

### Materials availability

This study did not generate new unique reagents.

## Data and code availability

The data that support the findings of this study are available from the lead contact upon request. The code for this study is available on GitHub: https://github.com/KaoutarAit01/axon-finder.git

## EXPERIMENTAL MODEL AND STUDY PARTICIPANT DETAILS

### Cyclic Immunofluorescence (CyCIF) Imaging

Prostate slides underwent CyCIF imaging using previously described methods ^36^, which involve repeated cycles of staining, imaging, and signal inactivation to achieve a high level of multiplexed protein imaging. 40 proteins were imaged, and the analysis for this paper is focused on two of these markers: TUBB3 ^37^ and NCAM1 ^38^. The process of primary antibody staining, imaging, and quenching was repeated for 10 rounds, using four distinct antibodies along with a DAPI nuclear stain in each round. A quenched signal was captured following rounds 3 and 10 and before the initial round of staining.

## METHOD DETAILS

### Patients Cohort

The study involved a group of 17 patients, aged at diagnosis between 46 and 81 years old. Based on the analysis of clinical data including PSA score, Gleason grades and lymph involvement, CAPRA-S risk scores were assigned to each patient ^31^. We grouped each patient as low-, intermediate- and high-risk based on the assigned CAPRA-S scores (Table 1).

**Table 1.** Summary of patients’ cohort.

|  |  | CAPRA-S |  |  |
| --- | --- | --- | --- | --- |
|  |  | Low-risk | Intermediate-risk | High-risk |
| PSA | 0-4 | 5 | 0 | 0 |
|  | 4-10 | 1 | 5 | 1 |
|  | > 10 | 0 | 2 | 2 |
| T stage | T2 | 7 | 4 | 1 |
|  | T3 | 0 | 2 | 3 |
|  | T4 | 0 | 0 | 0 |
| Primary Gleason grade | 3 | 7 | 0 | 0 |
|  | 4 | 0 | 5 | 4 |
|  | 5 | 0 | 1 | 0 |
| Secondary Gleason grade | 3 | 7 | 1 | 0 |
|  | 4 | 0 | 2 | 4 |
|  | 5 | 0 | 3 | 0 |
| Lymph node invasion | Yes | 6 | 1 | 0 |
|  | No | 0 | 2 | 4 |
| Recurrence | Yes | 0 | 0 | 1 |
|  | No | 6 | 5 | 2 |

### Image preprocessing and ground truth labeling

We generated a single image by registering and calculating the mean intensity values from TUBB3 and NCAM1 images. Subsequently, the histogram stretching step was implemented to improve the image quality for feature extraction. This technique effectively increased the image’s contrast and dynamic range, enhancing the visibility and accuracy of the markers’ distribution within the sample. The axons were delineated through a manual segmentation process, utilizing the Napari software. This procedure involved a hands-on, pixel-by-pixel delineation of the axonal structures within four whole-section images, ensuring that the segmentation task was executed with the utmost accuracy and attention to detail. A total of 997 axons were manually annotated across four slides. Our resulting single-channel combined images are 10000 pixels in size (width and height), which presents a challenge for processing with standard deep-learning algorithms because handling large images involves a trade-off between memory efficiency, computational speed, and maintaining relevant information for accurate model training and inference ^39^. To address this, we performed tiling to reduce the dimensionality of the data. Rather than passing the entire image as input, we divided it into smaller 15% overlapping images (1-channel tiles), each measuring 512 × 512 pixels. To enhance the representation of positive patches (i.e., regions positive for TUBB3 and NCAM1 staining), we augmented them by applying different rotations, flipping, sharpening, and shearing. This augmentation process led to a cumulative count of 11,680 crops that we split into 80% for training and 20% for validation of positive patches. Another dataset was created using only TUBB3 images as TUBB3 marks the majority of neurons whereas NCAM1 only marks a subset. We investigated whether using two markers (TUBB3 and NCAM1) improves the consistency of the axon’s segmentation using the same model. Upon the completion of this segmentation phase for each tile, we proceed to the reintegration stage where the segmented tiles are stitched, to reconstruct the segmentation map of the entire image.

### Deep Neural Network Architecture

A U-Net model was utilized to undertake the segmentation task. Our U-Net architecture is composed of the first four Blocks of the ResNet101 encoder network ^40^ responsible for capturing increasingly complex features from the input image. Following the encoder, there is a usual U-Net decoder that upscales these features to generate a segmentation map. Each decoder layer is structured with 3×3 convolutional operations, followed by a rectified linear unit activation and dropout regularization. The process of down-sampling is achieved through a 2×2 max-pooling operation. In the decoding phase, transpose convolutions are employed to gradually restore the spatial dimensions and reduce the number of channels in the feature maps to their original size. To enhance the learning process, skip connections are integrated at different scales. These connections concatenate the encoder’s output at each down-sampling layer with the corresponding up-sampling layer in the decoder. The primary objective of these skip connections is to propagate features from the same scale to each decoding layer, aiding in the understanding of both global and local contextual information. In the final up-sampling stage of the U-Net, a Softmax function is applied to convert the output into [0,1] scores, indicating the likelihood of each pixel belonging to one of two classes (i.e., axon vs. background).

### Training Procedure

Training the U-Net and optimizing its performance in our dataset involves experimenting with various hyperparameters. We systematically explored the impact of learning rates, batch sizes, and optimizer choices to enhance the model’s accuracy and efficiency.

We used different batch sizes for our model training setup (16, 32, 64, 128). Batch size influences the training speed and memory requirements, and its choice significantly affects the model convergence and generalization. Additionally, we assessed the performance across two distinct optimizers: Adaptive Movement Estimation algorithm (Adam) and Stochastic Gradient Descent (SGD). This optimizer governs the update rules for model parameters during training. Incorporating these variations in learning rate, batch size, and optimizer selection into our training process allowed us to systematically optimize our model for segmentation, achieving higher precision and robustness tailored to our specific requirements.

- Hardware: We opted for parallel training using a server with high computing power and Nvidia A40 GPUs which are high-performance GPUs designed for data-intensive tasks.
- Hyperparameter: We trained our model using these hyperparameter optimizers for Adam: learning rate (0.0001), batch size of (32), and epoch counts of (500).
- Training Period: ∼20 hours of intensive training duration. It underscores the substantial computational effort invested in refining the U-Net-ResNet-101 model parameters to enhance axon segmentation accuracy.

The training procedure was carried out twice, first by using a single marker (TUBB3) and second by using the combined image (TUBB3 and NCAM1).

### Evaluation Metrics

The AxonFinder segmentation method needs to assess both errors at the object level (Axon detection) and the pixel level (Axon shape and size). The F1-score is the employed metric for axon detection. In this context, it considers true positives (TP), which is the count of all ground truth objects associated with a segmented (detected) object false positive (FP), which is the count of all segmented objects without a corresponding ground truth object and false negatives (FN). The F1-score is defined as follows:

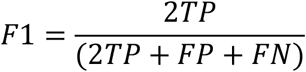

Furthermore, Intersection over Union (IoU) was also used to indicate the correlation between predicted pixel values and ground truth values. The IoU is defined as follows:

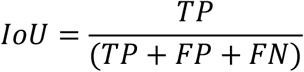

The weights of the best model in terms of the F1-score and the IoU were saved to infer it on the whole dataset.

## QUANTIFICATION AND STATISTICAL ANALYSIS

All procedures involving statistical analysis were described in the Method details section. The number of samples used in each test was described in the Results sections.

**Supplementary Figure 1. Spatial autocorrelation of axons based on TUBB3 and NCAM1 expression levels. (**A) Boxplot of TUBB3 expression levels stratified by CAPRA-S risk categories. (B) Spatial distribution of TUBB3 expression highlighting High (blue) vs. Low (orange) axons. (C) Ripley’s L function analysis for TUBB3 shows spatial clustering for Low TUBB3 axons (orange). (D) Boxplot of NCAM1 expression levels stratified by CAPRA-S risk categories. (E) Spatial distribution of NCAM1 expression highlighting High (blue) vs. Low (orange) axons. (C) Ripley’s L function analysis for NCAM1 shows spatial clustering for High NCAM1 axons (blue).

**Supplementary Figure 2. Morphological features of segmented axons do not show significant variability across the three patient groups. (**A) Samples from the dataset depicting large vs. small, branched vs. unbranched, and elongated vs. circular morphological features. (B) The distribution of the different axons’ morphological features for low-, intermediate- and high-risk patients. (C) Boxplot showing the morphological features stratified by CARPA-S risk categories.

**Supplementary Figure3**. GMM thresholding for the different axons’ features.

**Supplementary Figure 4**. The optimal number of k-means clusters is obtained when k=3.

