## Supplementary figures and images for "AxonFinder: Automated segmentation of tumor innervating neuronal fibers"

### Supplementary Figure 1

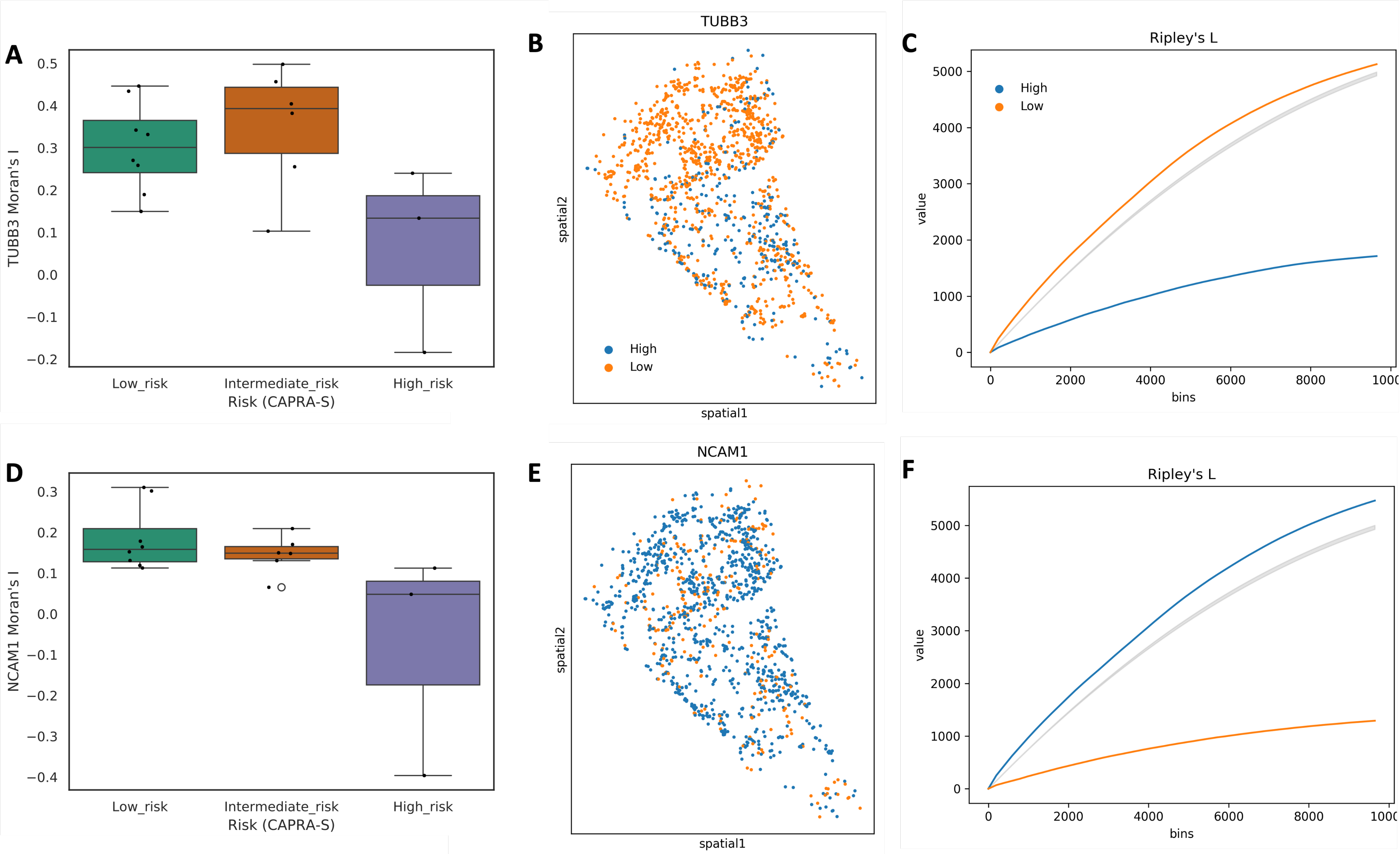

### Supplementary Figure 2

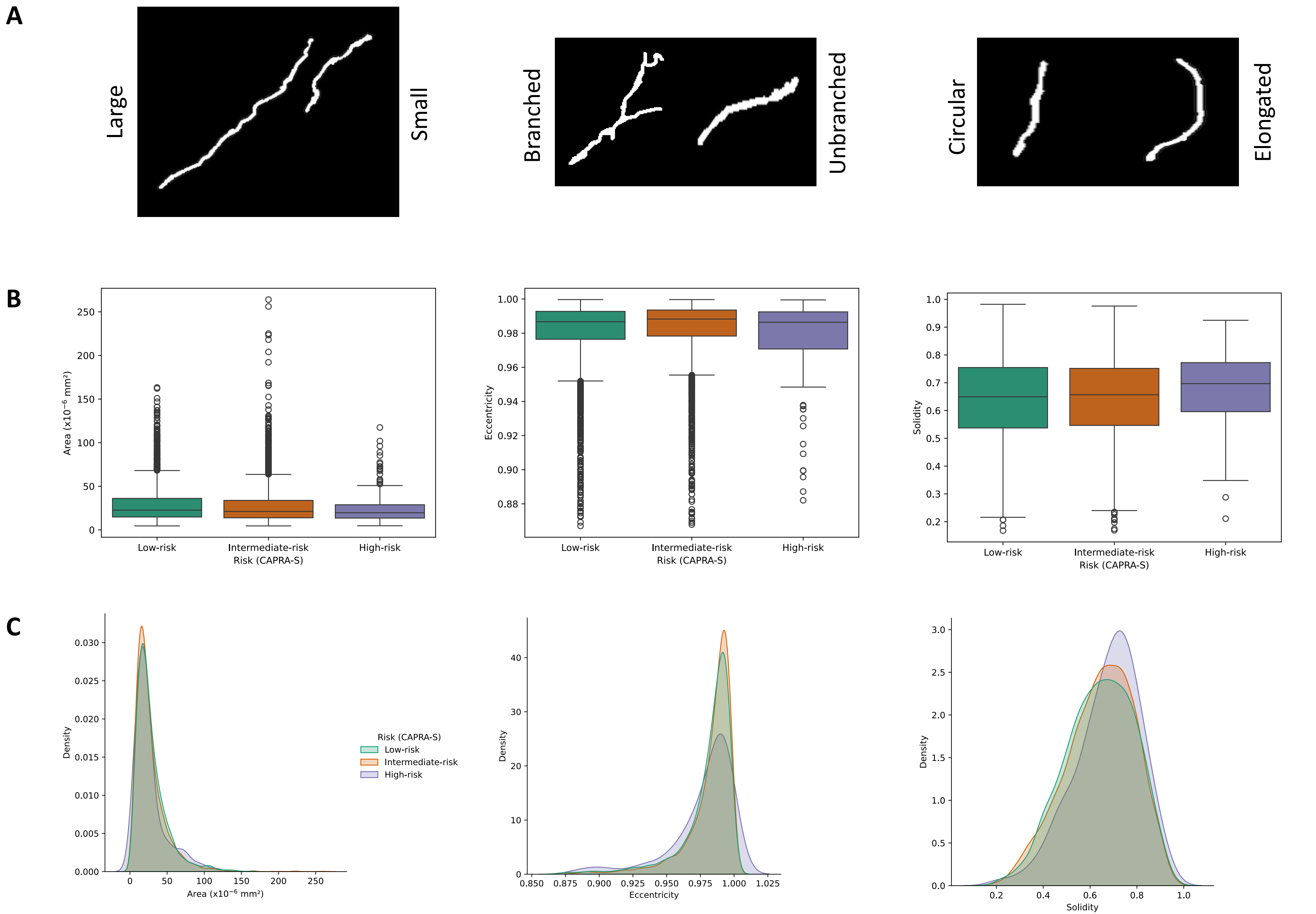

### Supplementary Figure 3

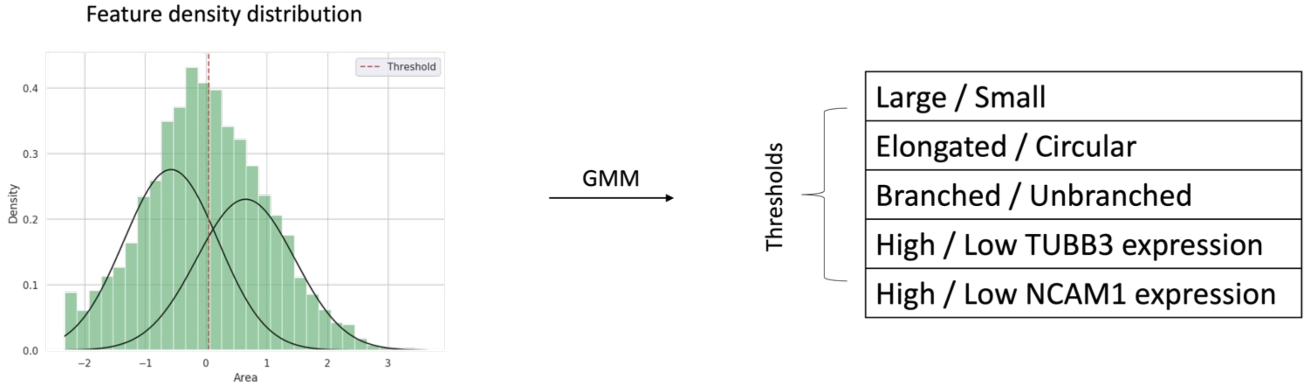

### Supplementary Figure 4

Elbow

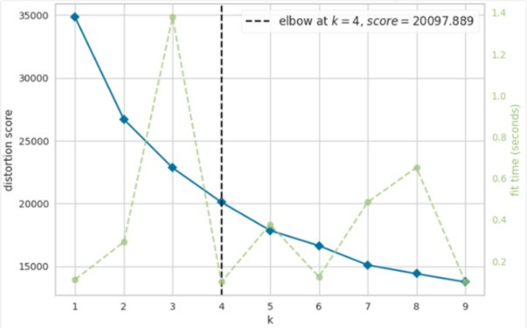


K-Means

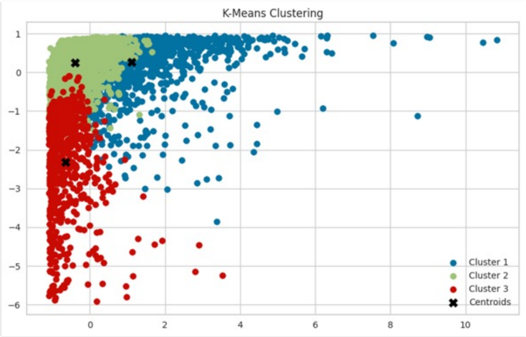
